# Multi-dimensional protein solubility optimization with an ultra-high-throughput microfluidic platform

**DOI:** 10.1101/2022.10.21.513267

**Authors:** Nadia A. Erkamp, Marc Oeller, Tomas Sneideris, Hannes Ausserwӧger, Aviad Levin, Timothy Welsh, Runzhang Qi, Daoyuan Qian, Hongjia Zhu, Pietro Sormanni, Michele Vendruscolo, Tuomas P.J. Knowles

## Abstract

Protein-based biologics are highly suitable for drug development, as they exhibit low toxicity and high specificity for their targets. However, for therapeutic applications, biologics must often be formulated to very high concentrations, making insufficient solubility a critical bottleneck in drug development pipelines. Here, we report an ultra-high-throughput microfluidic platform for protein solubility screening. In comparison with previous methods, this microfluidic platform can make, incubate, and measure samples in a few minutes, uses just 20 micrograms of protein (> 10-fold improvement) and yields 10,000 data points (1000-fold improvement). This allows quantitative comparison of formulation additives, such as salt, polysorbate, histidine, arginine and sucrose. Additionally, we can measure how solubility is affected by different concentrations of multiple additives, find a suitable pH for the formulation, and measure the impact of single mutations on solubility, thus enabling the screening of large libraries. By reducing material and time costs, this approach makes detailed multi-dimensional solubility optimization experiments possible, streamlining drug development and increasing our understanding of biotherapeutic solubility and the effects of excipients.

## Introduction

Proteins, peptides and antibodies, collectively known as biologics, have been the fastest growing drug class over the past decade.^1^ This ascent can be mainly attributed to their inherent low toxicity, high specificity, and favourable pharmacokinetics in comparison to small molecules.^2^ However, many such compounds lack sufficient solubility at the early stages of the drug development process, often resulting in costly late-stage failures, or yielding products that can only be administered intravenously, which usually requires long hospital visits for each dose.^3–6^ Proteins have evolved to be as soluble as necessary to sustain the required concentrations for their optimal biological functions.^7–9^ The in vivo concentrations of individual proteins or antibodies, however, are far below 100 mg/mL, which is typical in therapeutic formulations destined to subcutaneous injection.^10^

Solubility measurements remain material- and time-intensive, with the result that they are seldom incorporated at the early stages of drug discovery, where the number of candidates to screen is very high.^11^ The solubility of proteins can be measured *in vitro* using ultrafiltration and ultracentrifugation.^12,13^ Relative solubility measurements have also been developed to rank different proteins or formulation conditions with polyethylene glycol (PEG) or ammonium sulphate precipitation.^14–16^ This reduces material requirements and makes it easier to define solubility. Unlike compounds like salt, which in solution are either present as a solid or dissolved, proteins often populate plenty of different aggregated states, including small oligomeric species and ordered fibrils. This complexity makes the boundary between the soluble and insoluble phase ultimately arbitrary, and operationally dependent on the method used to separate the two (e.g., the speed and time of centrifugation).^7^ The relative solubility in these precipitation assays is defined as the amount of PEG or ammonium sulphate that is required for a protein to precipitate out of solution.^12–14^ These methods, however, are relatively low-throughput and still require significant amounts of purified protein material. Moreover, the impact of formulation parameters, such as pH, ionic strength, and additives typically used in biotherapeutic formulations are often only assessed by screening each parameter individually while keeping the others constant^17–20^, or only at the final-stages of preclinical development.^20–22^ It remains highly challenging to carry out multidimensional screenings of formulation parameters, in which solubility is measured by varying two or more parameters together (e.g., the pH and the concentrations of some excipients).^20,21,23^ Obtaining solubility data by sampling the multidimensional formulation space as exhaustively as possible is critical, as the solubility of biologics is highly formulation dependent.^20,21,23^ The solubility of proteins can be estimated *in silico* with methods such as CamSol^7,14^ and SAP^24^. However, such methods cannot be used for the exhaustive screening of formulation conditions, and it remains difficult to base critical decisions solely on computational predictions. Therefore, a new material- and time-efficient experimental method is required to screen candidates at the early stages of development, and to optimize protein solubility with respect to various variables, screened individually or in combination.^11^

Here, we present a microfluidic platform to measure the relative solubility of proteins at ultra-high-throughput and low cost. We demonstrate that this approach can create over 10,000 data points from only 20 μg of purified protein. In comparison with previous methods,^12–14^ this is a 1000-fold increase in datapoints and a greater than 10-fold decrease in required material. This ultra-high-throughput approach enables optimization of multiple variables in one experiment. In this work, we quantify how effective different formulation additives are at increasing solubility of a protein, measure solubility at different pH values, and select the protein variant with the highest solubility. Thus, this platform has the potential to streamline the drug development process and to greatly increase our understanding of the solubility of biotherapeutics and the effects of excipients.

## Results and discussion

### Measuring the relative solubility of biologics using microfluidics

We have capitalised on advancements in microfluidic technology^25,26^ to generate thousands of microdroplets containing protein, additives and precipitants (**Figure 1**, top). Each droplet gives an individual data point where the protein is in a different environment. Through modulating the flow rates of solutions containing protein, buffer, PEG and other compounds, droplets with an array of compositions are created. A fluorescent dye, either free in solution acting as a barcode or protein-bound, is added to the solutions. After incubating the droplets for 5 minutes, they are imaged at the wavelengths corresponding to the added dyes. The concentrations of the compounds in the droplet can then be inferred from fluorescence intensity. Moreover, as the protein is fluorescently tagged, we can directly observe if it is found homogenously in the droplet, or if it has formed precipitates, which appear as bright specs in the droplets. Droplet detection and analysis of their contents are performed by previously developed Python-based image analysis software^26^. By fitting data using a support-vector machine (SMV) algorithm (Methods, Supplementary Information), we can determine precipitation probability under different conditions. Next, we can construct phase diagrams, like a 2D diagram (**Figure 1**, bottom left), or determine the boundary at a certain protein concentration (**Figure 1**, bottom right). In these diagrams, the data points are shown, as well as the precipitation probability, with blue and red indicating a very low and very high chance, respectively, to form aggregates. The white area shows the phase boundary between the mixed phase and aggregated phase, and it is predicted based on the data. The thickness of this boundary indicates how accurately we have measured it and gives the standard deviation of the measurement. In **Figure 1**, we observe that the relative solubility of lysozyme is 5.7 ± 0.6% PEG at 1 mg/mL, consistent with previous measurements performed with a standard PEG-precipitation assay.^23^

**Figure 1.**
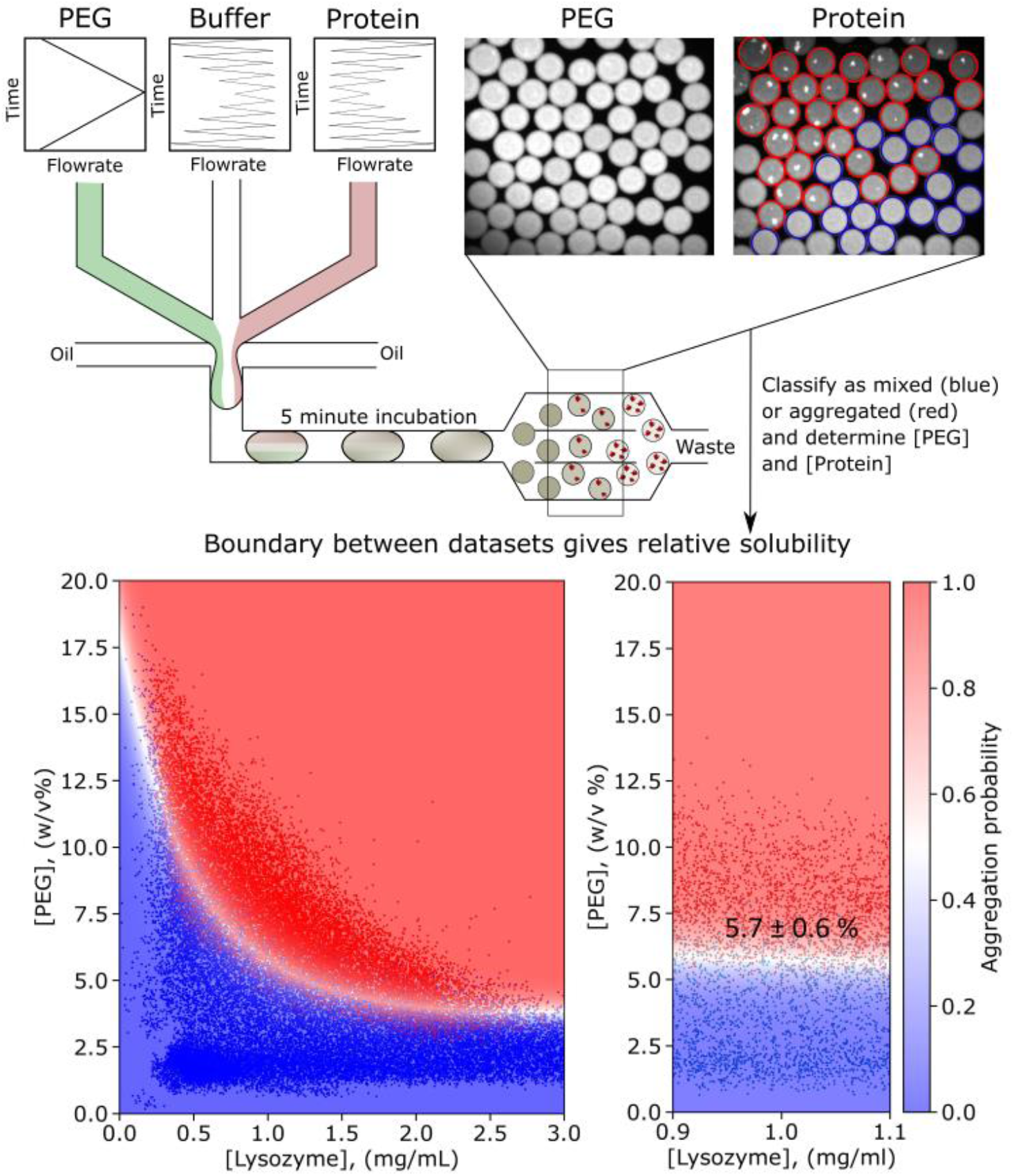
Setup for measuring protein solubility in an ultra-high-throughput aggregation assay. Water-in-oil droplets of ~100 pL are created by mixing solutions containing protein, buffer and PEG at various ratios. The droplets are incubated for 5 minutes and imaged at the wavelengths corresponding to the fluorescent dyes added solutions. A Python script^26^ is used to determine the concentration of each component and to classify them as containing a mixed solution (blue) or aggregates (red). We fit the data with an SVM algorithm and determine the aggregation probability. This procedure enables us to determine the phase boundary (white region) and the relative solubility value. In this example, we find that lysozyme has a relative solubility of 5.7 ± 0.6% PEG at pH = 7 at 1 mg/mL. The 2D graph on the left and the selection on the right displays 38,678 and 3,872 data points, respectively.

Using the microfluidic platform, the average volume per sample is just 100 pL. Droplets are created at a rate of 100 droplets/second and thus thousands of datapoints can be prepared in very minimal time. Due to the small size of samples, the incubation time is also greatly reduced, from hours or days^12,14^ to 5 minutes. We found that varying the incubation period from 2 minutes to 3 hours yielded no significant difference in the observed relative solubility, suggesting that our measurements are performed at near-equilibrium conditions (**Supplementary Figure 1**). Additionally, this method provides highly reproducible measurements (**Supplementary Figure 2**). Instead of PEG, ammonium sulphate is also often used to measure relative solubility. Ammonium sulphate can similarly be used in our setup for relative solubility measurements (**Supplementary Figure 3**).

### Optimization of formulations

To develop a protein drug, it is important to identify a formulation that enables a long shelf life.^27^ The compounds that are most often added to therapeutic formulations are sodium chloride (NaCl), histidine, arginine, sucrose and polysorbate 20 or 80.^28^ These compounds are added to reduce the viscosity (arginine), buffer the pH (histidine), provide a ionic osmotic pressure adjuster (NaCl), provide a non-ionic osmotic pressure adjuster (sucrose) and as a surfactant (polysorbate).^28^ In the absence of these compounds, the relative solubility of BSA at pH = 5 is 7.3 ± 0.7 w/v % PEG (**Supplementary Figure 2**). The addition of these compounds can significantly increase the relative solubility to up to 18 w/v% PEG (**Figure 2**). Moreover, with the microfluidic platform, we can quantitatively compare the ability of the additives to improve protein solubility. We find that, within the explored concentration range, the additives improve the solubility linearly, as would be expected in an ideal solution.^29^ By comparing the slope of the white boundary, we can rank these compounds according to their ability to increase protein solubility. The ranking, from least to most effective, is NaCl, histidine, arginine, sucrose and polysorbate 20 and 80. This matches expectations based on the roles of these compounds in the formulation, considering the surfactants increase the solubility most efficiently. Notably, while small amounts of NaCl improve the solubility at 0.020 PEG%/mM, adding additional NaCl beyond 350 mM only increases the solubility by 0.0023 PEG%/mM. Initially, in the salting-in regime, more salt interacts with the charges on the protein, reducing the inter-protein interactions and thus the propensity to form aggregates. However, adding further amounts of salt will not influence the solubility much, or could even harm the solubility in the salting-out regime.^16^ Using the measurements in **figure 2**, we can rationally design a formulation to improve solubility of the protein under scrutiny. For example, we may choose to use polysorbate 80 instead of polysorbate 20 and would prefer to add about 350 mM NaCl. Additionally, we would prefer to add more sucrose over arginine and histidine to improve the solubility.

**Figure 2.**
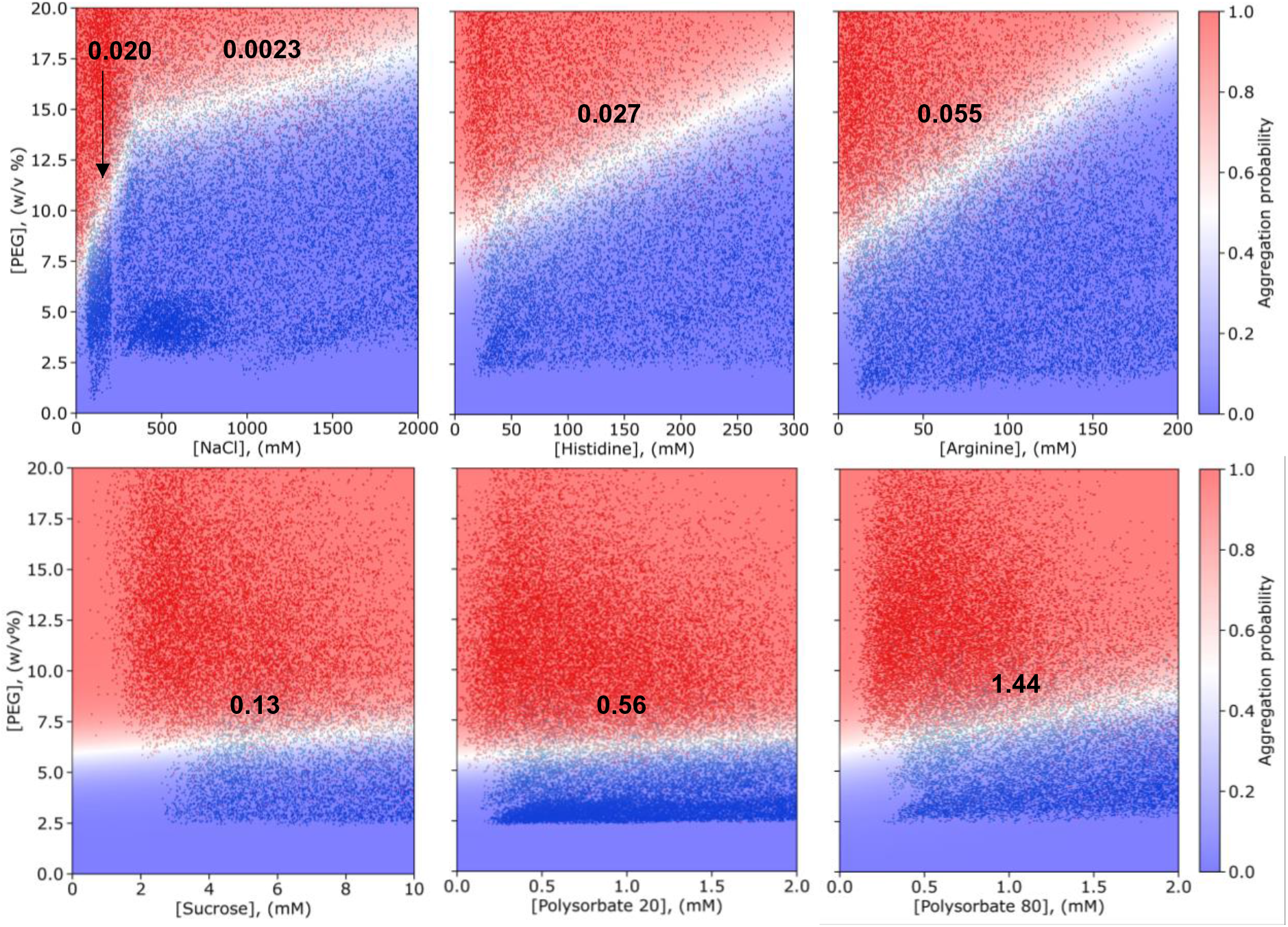
Formulation optimization screens with commonly used additives. Various amounts of additives are added to a 1 mg/mL BSA solution at pH = 5, increasing the relative solubility from 7.3 ± 0.7 w/v % PEG linearly in most cases (**Supplementary Figure 2**). The slope of the white boundary displays how much %PEG the relative solubility is increased per mM of additive. Thus, the ability of formulation additives to improve the solubility can be compared quantitatively. Of the 6 compounds tested, the surfactants polysorbate 20 and 80 most effectively improve solubility. The amount of data points shown is 18914, 20826, 15695, 23716, 35354 and 28750 for the graph of NaCl, histidine, arginine, sucrose, polysorbate 20 and 80, respectively.

### Directly comparing excipients

Using this platform, we can also compare different additives in one experiment, saving additional time and material when improving the formulation. Moreover, we can see how the solubility is affected by the presence of multiple additives. We varied the concentrations of 2 additives, NaCl and arginine, while keeping the amount of protein and PEG constant (**Figure 3a**). We obtain aggregates (red) in the absence of NaCl and arginine (bottom left corner), since we have 1 mg/mL BSA and PEG = 13.5, 14 or 14.5% in this experiment. By adding NaCl or arginine, we can improve the solubility and obtain a mixed solution (blue). From the shape of the slope, which is linear, we can also conclude that NaCl and arginine work together linearly to improve the solubility. Since the slope is negative, we can see that both compounds improve the solubility. If the value is −1, the compounds are equally well capable of improving the solubility. A value lower than −1 or higher than −1 indicates the compounds on the x-axis or y-axis, respectively, is more efficient at improving the solubility of the protein. Here, we find that the slope is around −3.4 and thus that

**Figure 3.**
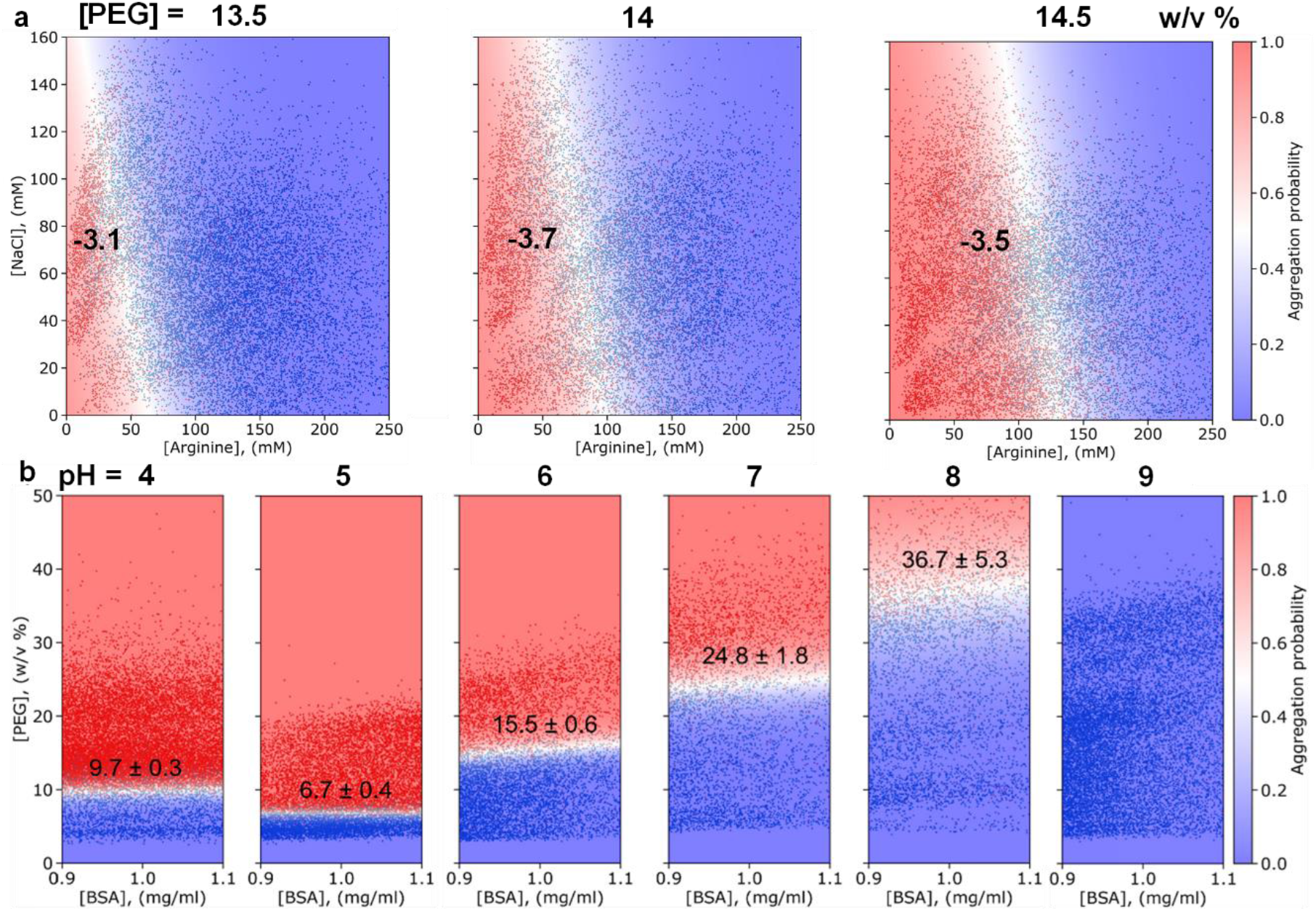
Comparison of formulation additives and optimization of the pH. **a** In the absence of NaCl and arginine, 1 mg/mL BSA at pH = 5 forms aggregates at PEG concentrations of 13.5, 14 and 14.5 w/v%. However, by adding NaCl and arginine, homogeneous solutions are obtained. The linear boundary shows that NaCl and arginine works together additively or linearly to increase the solubility. Based on the boundary slope, we can also conclude that arginine is more effective than NaCl at increasing the solubility. Adding more PEG decreases the solubility linearly, since it shifts the boundary to the right. From left to right, the amount of data points shown is 17143, 15780 and 16514. The slope, NaCl/arginine, is shown in the graph. **b** Relative solubility measurements of BSA at pH = 4, 5, 6, 7, 8, and 9, showing that this protein has the lowest solubility around pH = 5 and a higher solubility at a pH below or above 5. From left to right, the amount of data points shown is 16108, 10976, 7035, 4702, 4732, and 12423.

The compound on the x-axis, arginine, is more effective at improving the solubility. This matches with our finding in **Figure 2**, where arginine improved the solubility by 0.055%/mM and NaCl at these concentrations by 0.020%/mM. Three diagrams were produced at PEG = 13.5, 14 and 14.5%. We observe that the boundary shifts linearly to the right and that more NaCl and/or arginine are required to obtain a mixed solution. This is consistent with the fact that PEG promotes aggregation.

### Optimizing the pH of a formulation

pH is another important factor in protein solubility that can be scanned with the microfluidic platform. We measured the solubility of BSA in a formulation buffered at pH = 4, 5, 6, 7, 8 and 9 (**Figure 3b**). BSA has a relative solubility of 9.7 ± 0.3 % PEG at pH = 4, at a solubility of 6.7 ± 0.4 % at pH = 5. Thus, in a formulation, pH = 4 would be more suitable. Alternatively, increasing the pH can improve the solubility significantly, with the solubility at pH = 9 being so high that the boundary could not be determined under these conditions. The measured values and trend compare well to previously reported measurements carried out with standard PEG-precipitation assays (**Supplementary Figure 4**).^23^ Notably, in comparison to the previous measurements,^23^ the microfluidic-based measurements shown in **Figure 3** require 90% less of the protein, have an incubation time of 5 minutes instead of 2 days, have a smaller measurement error as they comprise thousands of data points instead of just tens.

### Comparison of the solubility of different mutational variants

Another way to improve protein solubility is to mutate the protein to identify more soluble variants, while maintaining its function^7,14^. Moreover, screening campaigns of biologics typically yield hundreds of candidates that may differ by as little as one mutation.^2,30,31^ Our microfluidic platform can be employed to measure the solubility of protein mutants. **Figure 4a** shows the structure of IgG4 antibody, of which the wild type and 6 of its previously designed mutational variants were used.^15^ The CamSol method was used to estimate the relative solubility of these variants,^15^ showing that variants 1-3 are expected to be less soluble than the wildtype, while variants 4-6 are expected to be more soluble (**Figure 4b**). **Figure 4c** shows the solubility measurements of the mutants and wildtype containing over 1500 data points each. The wildtype has a relative solubility of 11.1 ± 1.1 % PEG. The variants 1-3 indeed have a lower relative solubility and variants 4-6 a higher solubility. Comparing these measured values with the prediction by CamSol^15^, we observe that the variants with the lowest CamSol score, meaning they are predicted to aggregate easiest, indeed require the least amount of PEG to form aggregates. Even if these variants differed only by 2 to 4 point mutations in the context of a full IgG of about 1350 residues, the microfluidic platform allows us to select variant 6 as the one that is most soluble and thus optimize the protein sequence.

**Figure 4.**
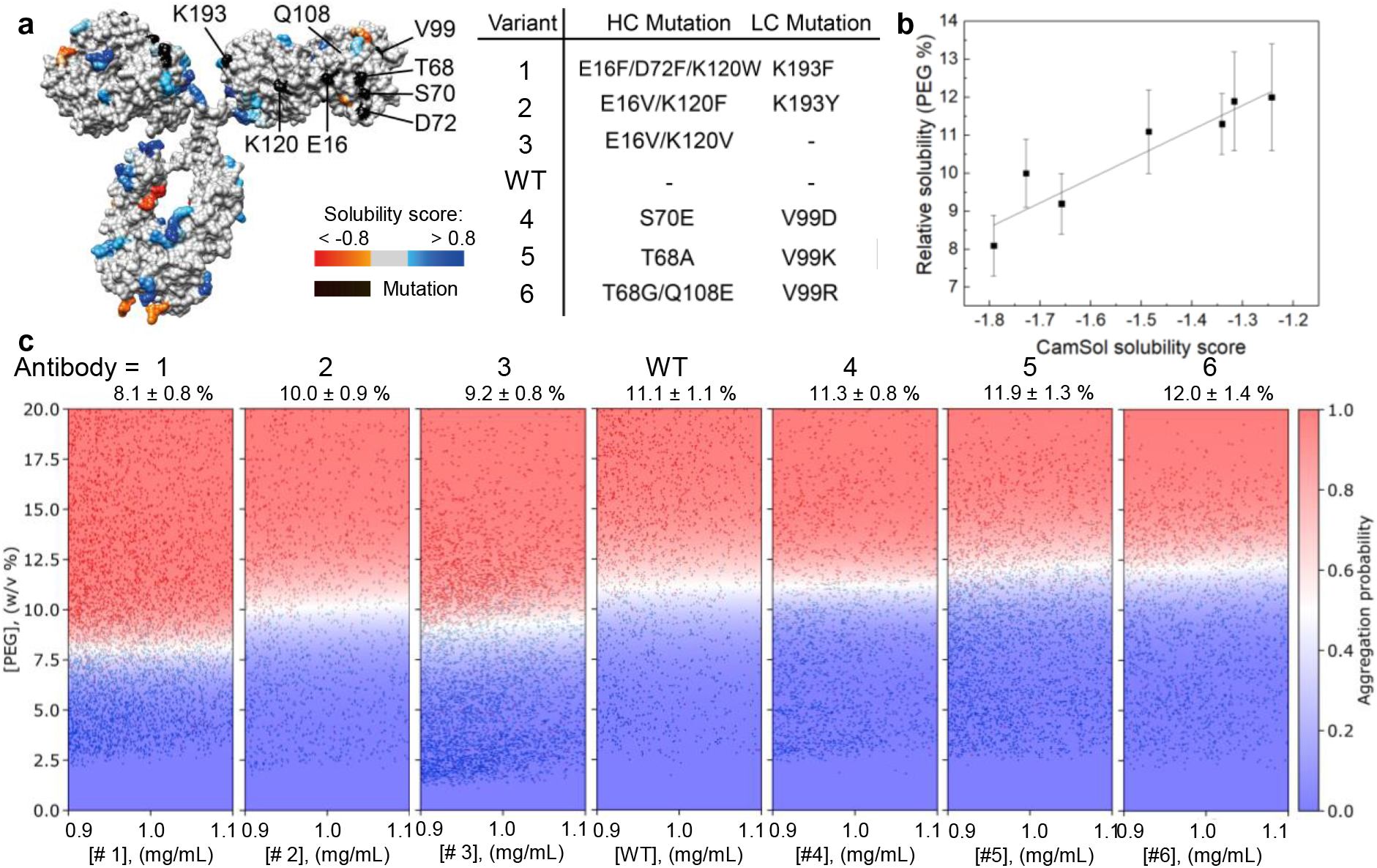
Solubility measurements of an IgG4 antibody and 6 mutational variants. (**a**) Structure of the IgG4 antibody, with the solubility score of regions and locations of mutations depicted on the surface. 6 variants are used containing 2 to 4 point mutations. (**b**) The relative solubility of the 6 variants and the wildtype is measured by our microfluidic platform and plotted against the CamSol solubility score.^15^ A clear trend in solubility between these proteins can be measured, despite the small difference in sequence. r^2^ = 0.86. (**c**) Microdroplet-based relative solubility measurements of the variants and wildtype. From left to right, the diagrams contain 5316, 1733, 5308, 1899, 2360, 3605 and 2326 data points.

## Conclusions

Optimizing protein solubility remains a significant bottleneck in the development of protein-based drugs. High solubility is required for long-term storage and to ensure efficient administration. However, available techniques have high material requirements and not enough throughput to measure protein solubility during early development or to optimize the formulation extensively. We have presented a microfluidic platform that addresses both concerns. We can quantify the solubility increase obtained by different kinds and different concentrations of formulation additives, can optimize the pH and study how the additives influence solubility in each other’s presence. The relative solubility can be determined with either PEG or ammonium sulphate, both industry standards. Additionally, our method enables the accurate solubility ranking of protein variants, even when these differ only by a few point mutations. Our microfluidic platform provides a highly quantitative strategy for improving a wide range of aspects influencing protein solubility and can aid in the development of new protein-based drugs.

## Supporting information

Supplementary Information: Supplementary figures 1-4 and materials and method section

## Acknowledgements

We thank Nikolai Lorenzen and Gaetano Invernizzi (Novo Nosdisk A/S) for their feedback. We thank Bjarne Gram Hansen, Birgitte Friedrichsen and Anne Lee Andersen (Novo Nordisk A/S) for their efforts in organizing plasmid generation and antibody expressions. The research leading to these results has received funding from a Royall Scholarschip (N.A.E.), AstraZeneca (M.O.), the European Union’s 279 Horizon 2020 research and innovation programme under the Marie Skłodowska-Curie grant MicroREvolution 280 (agreement no. 101023060; T.S.), the Newman Foundation (T.S., T.P.J.K.), Global Research Technologies Novo Nordisk A/S (H.A, T.P.J.K.), the Harding Distinguished Postgraduate Scholar Programme (T.J.W.), a Krishnan-Ang Studentship (R.Q.), Trinity College (Cambridge Honorary Trinity-Henry Barlow Scholarship; R.Q.), the China Scholarship Council (H. Z.), a Royal Society University Research Fellowship (P.S., URF\R1\201461), Wellcome Trust Collaborative 283 Award 203249/Z/16/Z (T.P.J.K.) and the European Research Council under the European Union’s 285 Seventh Framework Programme through the ERC grants PhysProt (T.P.J.K., agreement no. 337969; 286)

## Author Contributions

N.A.E., A.L. and T.P.J.K. conceived the study. N.A.E., M.O., T.S., H.A., T.W., R.Q., D.Q., H.Z. and P.S. have performed investigation. P.S. M.V. and T.P.J.K. acquired funding. N.A.E., M.O. and T.P.J.K. wrote the original draft, all authors reviewed and edited the paper.

## Additional information

Supplementary information is available online for this paper. Data generated in the study are available on reasonable request from the corresponding author.

