## Supplementary Information: Supplementary figures 1-4 and materials and method section for "Multi-dimensional protein solubility optimization with an ultra-high-throughput microfluidic platform"

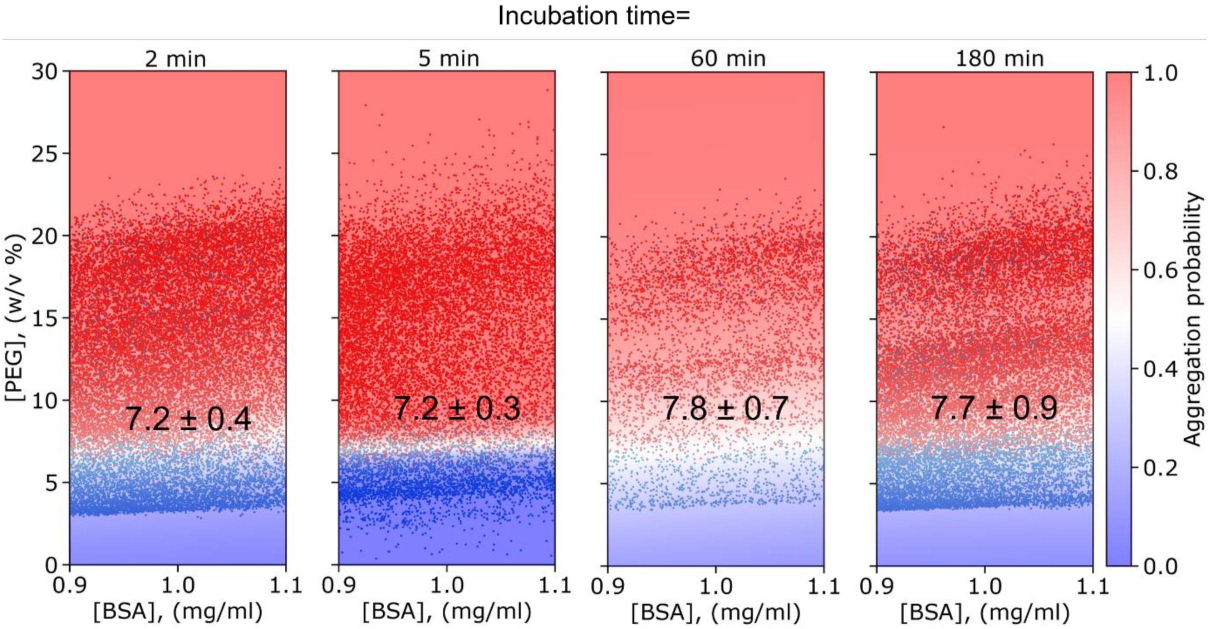


**Supplementary Figure 1. Relative solubility measurements after different incubation times.** To assess the effects of different incubation times, we measure the relative BSA solubility at pH = 5 following microfluidic droplets incubation for 2, 5, 60 and 180 min. Notably, while previous work required incubation times of up to 48 hours,[1] our smaller samples are mixed and can aggregate in much less time. All other data in this paper were acquired after a 5 min incubation period. From left to right, the graphs contain 21390, 15242, 4786 and 19846 data points.

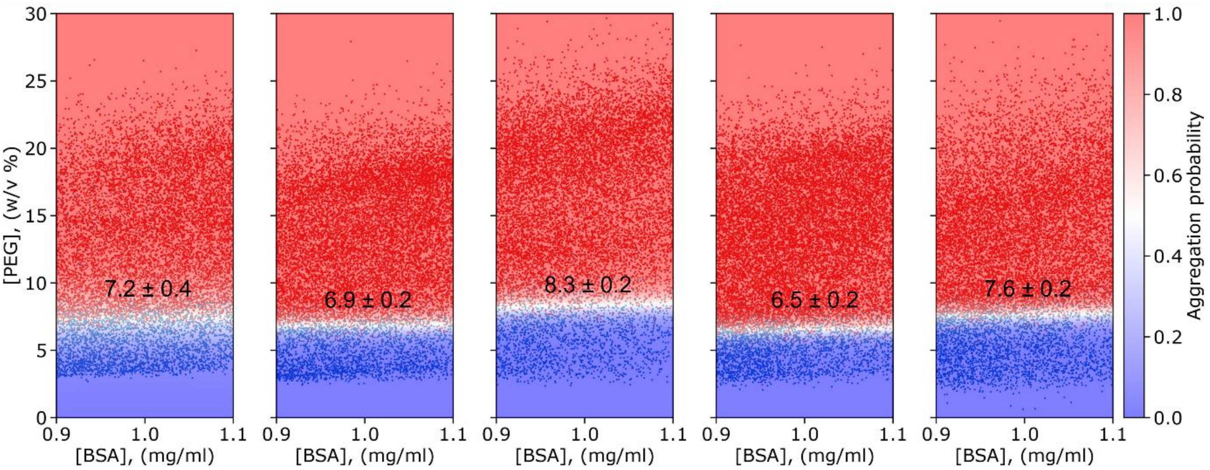


**Supplementary Figure 2. Robustness of the protein solubility measurement.** Relative solubility of BSA batches at pH = 5 was measured in different microfluidic devices on different days. The results are highly reproducible. Combining these 5 measurements, we also find that the relatively solubility of BSA at pH = 5 is 7.3 ± 0.7 w/v % PEG. From left to right, the graphs contain 12109, 14348, 12758, 17080 and 13481 data points.

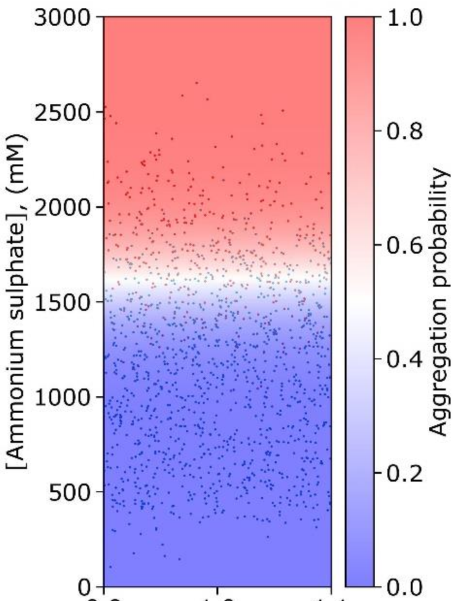


**Supplementary Figure 3. Relative solubility measurement of BSA with ammonium sulphate.** Both PEG and ammonium sulphate[2] are industry standards to determine the relative protein solubility. Here, we measure the relative BSA solubility at pH = 5 using ammonium sulphate, instead of PEG and find a relative solubility of 1600 ± 220 mM ammonium sulphate. 1437 data points are shown.

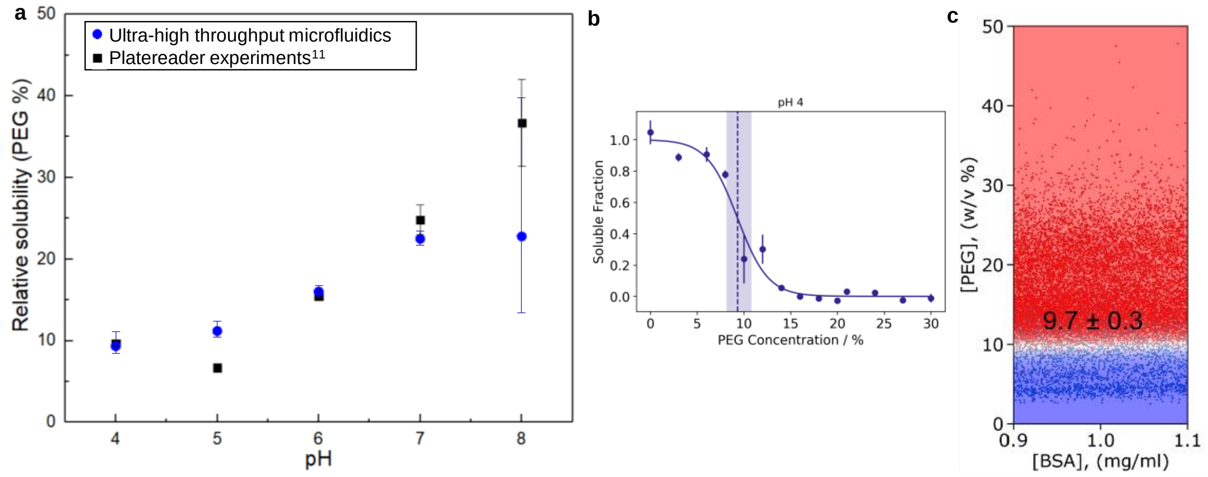


**Supplementary Figure 4. Comparison of measurements carried out on the microfluidic platform and on a plate reader. (a)** The relative BSA solubility at pH = 4, 5, 6, 7 and 8 is measured using out ultra-high-throughput microfluidic platform and previously determined using a platereader.[3] We see that we obtain a very similar trend and values. Notably, in comparison with the plate reader experiment, the microfluidic platform has an incubation time of 5 minutes instead of 48 hours, uses only 10% of the protein, results in a smaller measurement error, gives thousands instead of 15 data points and can screen for another variable, like protein concentration, at the same time. Values from **Figure 3** and a previous publication.[3] **(b)** Example of measuring relative solubility using a plate reader.[1] Briefly, BSA at pH = 4 is mixed to obtain a final concentration of 1 mg/mL with 15 different amounts of PEG using a pipetting robot and incubated at 4 °C. After 48 hours, the plate is centrifuged and the supernatant is transferred into a fresh plate. The amount of BSA that is dissolved in solution is compared with the total amount to give the soluble fraction. Via fitting, the amount of PEG at which 50% of the protein is in solution is determined, which is used as the relative solubility. The relative solubility was found to be 9.3 with an asymmetric error from 8.1 to 10.8.[3] **(c)** Measurement of BSA solubility at pH =4 using the ultra-high-throughput microfluidic setup, see also **Figure 3b**. The relative solubility was found to be 9.7 ± 0.3 using 16108 data points.

### Materials and Methods

### Materials

PEG (average molecular weight of 8000 g/mol), trichloro(1H,1H,2H,2H-perfluorooctyl)silane, HEPES, ammonium sulphate, NaCl, sucrose, arginine, histidine and polysorbate 20 and 80 were obtained from Sigma Aldrich. Alexa Fluor™ 488 Carboxylic Acid and Alexa Fluor™ 647 NHS ester was obtained from Thermo Fisher. Sylgard 184 Elastomer base and curing agent were obtained from Dow Corning Corporation. 24x60 mm No.1.5 glass slides were obtained from DWK Life Sciences. HFE-7500 was purchased from Fluorochem and fluorosurfactant was obtained from RAN Biotechnologies.

**Antibody expression and purification**

The antibodies were kindly provided by Novo Nordisk and were expressed and purified as reported previously[4]. Briefly, vectors with mutations were produced using pNNC340 (expression vector harbouring WT HzANTP heavy chain) and pNNC341 (expression vector harbouring WT HzANTP light chain) using QuickChange Lightning multi-site-directed and sitedirected mutagenesis kits, respectively (Agilent Technologies). Strings with the mutations (Thermo Fisher Scientific) with 15 basepair long overhangs on both sides were added to the PCR mix (KOD xtreme kit, Merck Millipore) with pNNC340-41 as template (InFusion HD cloning kit (Takara Bio). The material was transformed into E. coli DH5alpha competent cells (Thermo Scientific), and plated on LuriaBertani (LB) agar plates with carbencillin and incubated at 37°C overnight. Single colonies were inoculated in 2 mL of LB medium with carbenicillin and grown at 250 rpm shaking at 37°C overnight. A QIAGEN Plasmid Plus 96 BioRobot Kit (Qiagen) and a Biomek FXP pipetting robot (Beckman Coulter, Brea, US) was used to harvest the plasmids and the sequence was confirmed Eurofins Scientific's sequencing service. Clones harbouring the correct vectors were similarly to before re-transformed, plated, and single colonies were grown. Using GenElute HP maxiprep kit (Sigma-Aldrich), over 1 mg of each vector was obtained and transfected in a 1:1 ratio for the heavy and light chain into Expi293F™ cells (Thermo Fisher Scientific, Waltman, US) with a density of $3\partial 106$ cells $mL-1$ and over 95% viability (NC-3000 NucleoCounter (Chemometec). Cultures were grown at 36.5°C, 8% $CO_2$ and 125 rpm shaking. After 5 days, the cultures were harvested. The supernatant was filtered and the amount of antibody was quantified by Dip and ReadTM Protein A (ProA) Biosensors in an Octet system (Pal ForteBio). The protein was purified on an Äkta Express chromatography system using affinity purification using Mabselect Sure Protein A resin and then Superdex200 resin (GE Healthcare). The column was washed with 0.1 M HEPES pH 7.4, 150 mM NaCl, and antibodies were eluted with 0.1 M sodium formate, pH 3.5 into a pre-equilibrated gel filtration column, with a running buffer of 20 mM HEPES, 0.15 M NaCl, pH 7.4. Eluted fractions were collected in a 96-well plate. Fractions were pooled to obtain high purities and minimum higher molecular weight protein. The concentration of the antibody was determined by absorbance at 280 nm absorbance with Dropsense 96 (Trinean).

**Fabrication of microfluidic devices**

Standard lithography techniques were used to fabricate microfluidic devices.[5] Briefly, the device was designed in AutoCAD (AutoDesk) and printed on a photomask (Micro Lithography). UV expose was used to place the pattern of the mask on a silicon wafer coated with a 50 µm thick layer of SU8-3050 photoresist (Microchem), after heating at 95 °C for 45 minutes. Excessive SU-8 photoresist was removed using propylene glycol methyl ether acetate (Sigma), after heating 5 minutes at 95 °C. The wafer with SU-8 patterns, or master, was dried and served as a mould to make PDMS, poly(dimethylsiloxane), devices. PDMS base and curing agent were mixed in a ratio of 10:1 and baked for 1 hour at 65 °C. The PDMS was then removed from the mould and cleaned by sonication in ethanol. Holes were punched where liquids were put into or would go out of the device with a biopsy puncher. Both the PDMS and a glass slide were activated in an oxygen plasma oven (30 s, 40% power, Femto, Diener Electronics) before bonding together. The device was heated at 95 °C for 3 minutes. The channels were treated with 1% (v/v) trichloro(1H,1H,2H,2H-perfluorooctyl)silane in HFE-7500 (Fluorochem) for 1 minute and subsequently dried with nitrogen gas and then heated at 95 °C for 10 minutes.

**Microdroplet preparation and imaging**

The solutions to be combined to make microdroplets were prepared. In a typical experiment, a solution with just buffer (for example, 10 mM phosphate citrate at a certain pH value), a solution with protein mixed with 10 molar-% Alexa Fluor™ 647 NHS ester and left for 15 minutes in buffer, and a solution with 50 w/v % PEG 8000 in buffer with 2 μM non-reactive Alexa Fluor™ 488 carboxylic acid would be prepared. Depending on the amount of datapoints desired, between 5 and 50 μL each solution is loaded into the tubing (PTFE, 0.012"ID x 0.030"OD, Cole-Parmer). These solutions are pushed into the microfluidic device using gas-tight 0.1 mL syringes (Hamilton 1710) operated by syringe pumps (neMESYS modules, Cetoni). Additionally, a gastight 1 mL syringe (Hamilton 1710) is used to push HFE-7500 oil with 1.5% fluorosurfactant into the microfluidic device. The aqueous flowrates changed relative to each other, depending on the preferred concentrations in the droplets, with the sum of these flowrates being constant. The oil flow rate was between 50 and 200 μL/h, such that droplets of around 100 pL in volume were obtained. By flowing the solutions, the droplets are made at around 100 droplets/seconds. After 5 minutes of incubating the droplets, by making them travel through the device, they are pictured in all relevant wavelengths, typically at 488, 546 and 647 nm. Each set of images contains around 100 droplets and thus 100 datapoints. During the experiment shown in Supplementary Figure 1, the incubation time was varied by increasing the flow rate or stopping the flow for an extended amount of time, meaning the droplets were incubated for a longer or shorter amount of time.

To determine the compound concentrations based on the intensity of the pictures, a few calibration images were obtained. Droplets containing only 1 of the solutions, for each solution, were imaged as a reference. Additionally, pictures of a homogenous solution with the fluorescent dyes were taken to correct for illumination differences by the laser light in different spots. Lastly, a background image was taken, without any sample or any light on, to observe the background noise the camera will always pick up. For all these calibration images, 10 images were taken and averaged.

**Image analysis**

The images of the droplets and calibration images were processed by a previously published Python script[6] that analyses microfluidic droplets, obtains concentrations based on intensities and looks for inhomogeneous intensities within droplets, which is how aggregates are detected. Further analysis of pictures was performed using Fiji. Figure 2a was created using Chimera.

**Data fitting**

The phase boundary between mixed and aggregate-containing droplet populations was determined by fitting the data using a support-vector machine (SVM) algorithm with linear or 2$^{nd}$ degree polynomial kernel, programmed in Python.
